# CEP290 myosin-tail homology domain is essential for protein confinement between inner and outer segments in photoreceptors

**DOI:** 10.1101/660738

**Authors:** Poppy Datta, Brandon Hendrickson, Sarah Brendalen, Avri Ruffcorn, Seongjin Seo

**Affiliations:** Department of Ophthalmology and Visual Sciences, University of Iowa College of Medicine, Iowa City, IA 52242, USA; Institute for Vision Research, University of Iowa, Iowa City, IA 52242, USA

**Author notes:** Correspondence to: Seongjin Seo, PhD, 375 Newton Rd., Department of Ophthalmology and Visual Sciences, University of Iowa, Iowa City, IA 52242, USA.

## Abstract

Mutations in *CEP290* cause various ciliopathies involving retinal degeneration. CEP290 proteins localize to the ciliary transition zone and are thought to act as a gatekeeper that controls ciliary protein trafficking. However, precise roles of CEP290 in photoreceptors and pathomechanisms of retinal degeneration in *CEP290*-associated ciliopathies are not sufficiently understood. Using *Cep290* conditional mutant mice, in which the C-terminal myosin-tail homology domain is disrupted after the connecting cilium is assembled, we show that CEP290, more specifically the myosin-tail homology domain of CEP290, is essential for protein confinement between the inner and the outer segments. Inner segment plasma membrane proteins including STX3, SNAP25, and IMPG2 rapidly accumulate in the outer segment upon disruption of the myosin-tail homology domain. In contrast, localization of endomembrane proteins is not altered. Trafficking and confinement of most outer segment-resident proteins appear to be unaffected or only minimally affected in this mouse model. One notable exception is RHO, which exhibits severe mislocalization to inner segments from the initial stage of degeneration. Similar mislocalization phenotypes were observed in *rd16* mice. These results suggest that failure of protein confinement at the connecting cilium and consequent accumulation of inner segment membrane proteins in the outer segment combined with insufficient RHO delivery is part of the disease mechanisms that cause retinal degeneration in *CEP290*-associated ciliopathies. Our study provides insights into the pathomechanisms of retinal degenerations associated with compromised ciliary gates.

## INTRODUCTION

Photoreceptor cells in the retina are highly compartmentalized neurons. While proteins involved in the phototransduction cascade are confined to a compartment called the outer segment, ones responsible for energy metabolism and protein/lipid synthesis are confined to another compartment, the inner segment. Proteins that constitute the outer segment are synthesized in the inner segment and transported to the outer segment through a narrow channel, called the connecting cilium. The outer segment is a primary cilium-related compartment and the connecting cilium is equivalent to the transition zone in primary cilia. Since connecting cilia are the only conduit between the inner and the outer segments, they are situated at a critical place to control protein movement between the two compartments and maintain compartment-specific protein localization.

Although outer segments share many features with primary cilia, they have several unique features that are not observed in primary cilia (reviewed in [1]). For example, outer segments are much larger than primary cilia, accounting for 25-60% of the total cell volume depending on cell types (rod vs. cone) and species. Outer segments are also filled with membranous structures called discs. The large size and high membrane content of the outer segment provides a space to accommodate a large amount of transmembrane and lipidated proteins that constitute the phototransduction cascade. In addition, outer segments are constantly and rapidly regenerated: old discs are shed from the distal end of the outer segment and new discs are formed at the base [2]. Constant and rapid regeneration of a large organelle necessitates a high volume of lipid and protein transport through the connecting cilium. At the same time, proteins that are not authorized to pass the connecting cilium (in either direction) should be kept in their designated compartments. Several proteins at the connecting cilium are thought to organize a gate to control protein movement between the inner and the outer segments [3].

CEP290 is one of such proteins. In model organisms *Chlamydomonas reinhardtii* and *Caenorhabditis elegans*, CEP290 is found at the ciliary transition zone and required to build Y-shaped linkers that extend from the axonemal microtubules to the ciliary membrane [4–6]. Although cilia/flagella still form in the absence of CEP290 in these organisms, ciliary protein compositions are altered [4, 6]. In mammalian primary cilia, CEP290 is localized to the transition zone [7–9], and loss of CEP290 reduces ARL13B and ADCY3 levels within cilia while increasing the rate of ciliary entry of SMO in fibroblasts [10]. These studies establish the current model for CEP290 function: a ciliary gatekeeper that regulates protein trafficking in and out of the ciliary compartment at the transition zone [4–7, 10–13]. In photoreceptors, CEP290 is localized to the connecting cilium [14, 15] and expected to control protein movement between the inner and the outer segments.

Mutations in human *CEP290* cause various ciliopathies ranging from isolated retinal dystrophy (e.g. Leber congenital amaurosis (LCA)) to syndromic diseases such as neonatal lethal Meckel-Gruber Syndrome (MKS) with multi-organ malformations [16–22]. Despite considerable variations in phenotypic severity, retinopathy is present in almost all cases regardless of the involvement of other organs. This suggests that photoreceptors are particularly susceptible to deficiencies in CEP290 function. Based on the ciliary gatekeeper model, anticipated functions of CEP290 in photoreceptors include i) permitting or facilitating entry of outer segment-bound proteins into the outer segment, ii) blocking unauthorized entry of inner segment proteins into the outer segment, and iii) preventing diffusion of outer segment proteins into the inner segment. In line with the first function listed above, *Cep290* mutant mice fail to develop outer segments. The connecting cilium and the outer segment are entirely absent in *Cep290* null mice [15]. Partial loss of CEP290 function in *rd16* mice, which have an in-frame deletion of exons 36-40 (based on Reference Sequence transcript NM_146009), allows formation of membrane-bound connecting cilia, but outer segments are rudimentary and severely malformed [14, 15, 23]. These studies show that CEP290 is essential for outer segment biogenesis and, either directly or indirectly, required for the trafficking of outer segment proteins.

However, precise roles of CEP290 in photoreceptors and disease mechanisms that induce retinal degeneration in *CEP290*-associated ciliopathies are not sufficiently understood. This is partly because of the critical requirement of CEP290 for the outer segment biogenesis in mouse models. As described above, *Cep290* mutant mice have no or only rudimentary outer segments [14, 15, 23]. While these phenotypes demonstrate the requirement of CEP290 for the outer segment biogenesis, complete lack or severe malformation of the outer segment precludes further investigation of CEP290’s role in protein trafficking and confinement between the inner and the outer segments. For instance, although CEP290 is likely required for the trafficking of at least certain outer segment proteins, specifically which proteins require CEP290 is unclear. Requirement of CEP290 for protein confinement between the inner and the outer segments has not been demonstrated either. Contrary to the findings in mouse models, Cep290 (or the C-terminal half of Cep290) does not appear to be essential for outer segment biogenesis in zebrafish [24]. In *cep290*^*fh297/fh297*^ mutants, which have a nonsense mutation (p.Gln1217X) near the middle of the reading frame, retinas develop normally during embryogenesis. In addition, retinal degeneration is slow and limited to cones in this model. Interestingly, although RHO mislocalization is detected at 6 months post fertilization, degeneration of rods is not observed. Apart from RHO mislocalization, obvious signs of disrupted ciliary trafficking (e.g. accumulation of vesicular materials near the ciliary base in electron microscopy) are also not observed in this model. Therefore, precise roles of CEP290 including its gating functions remain to be determined in photoreceptors.

In this work, we sought to test the current model of CEP290 function as a ciliary gatekeeper in photoreceptors and advance our understanding of the pathomechanisms underlying *CEP290*-associated retinopathies. We avoided the aforementioned limitations of the *CEP290* mouse models with constitutive mutations by using a model with a conditional allele and disrupting CEP290 functions after the connecting cilium assembly. In addition, we reasoned that disruption of ciliary gates would have a profound impact on the localization of inner segment membrane proteins, because of the physical properties of the outer segment (i.e. large size, high membrane content, and continuous disc renewal) and its tendency to house membrane proteins [1, 25]. Therefore, we examined (mis)trafficking of not only outer segment-resident proteins but also various inner segment membrane proteins.

## RESULTS

### Characterization of the *Cep290*^*fl*^ conditional allele

To avoid the outer segment biogenesis defects in constitutive *Cep290* mutant mice, we employed a mouse line with a *Cep290* conditional allele (*Cep290*^*fl*^) [26] and aimed to disrupt CEP290 functions after the connecting cilium is assembled. The *Cep290*^*fl*^ allele has LoxP sites flanking exons 37 and 38 [26] (**Figure 1A**). Hereafter, we use the term *Cep290*^*Δ*^ to denote the *Cep290*^*fl*^ allele lacking these two exons after CRE-mediated excision. Removal of these exons causes a frameshift followed by a premature termination codon. This change may induce nonsense-mediated decay of the mutant mRNAs, rendering *Cep290*^*Δ*^ a null or a strong hypomorphic allele. If not, mutant mRNAs are expected to produce a C-terminally truncated protein (p.Leu1673HisfsTer6) with a predicted molecular weight of ∼196 kDa. This truncated protein is likely to maintain some functions of full-length CEP290. *rd1* (in *Pde6b*) and *rd8* (in *Crb1*) mutations present in the cryo-recovered mice were removed by breeding with wild-type 129S6/SvEvTac mice. For robust and consistent excision in the vast majority of photoreceptors after the connecting cilium assembly, *Cep290*^*fl/fl*^ mice were crossed to *iCre75* (*Cre* hereafter) transgenic mice that express CRE recombinase under the control of mouse *rhodopsin* (*Rho*) promoter in rods [27].

**Figure 1.**
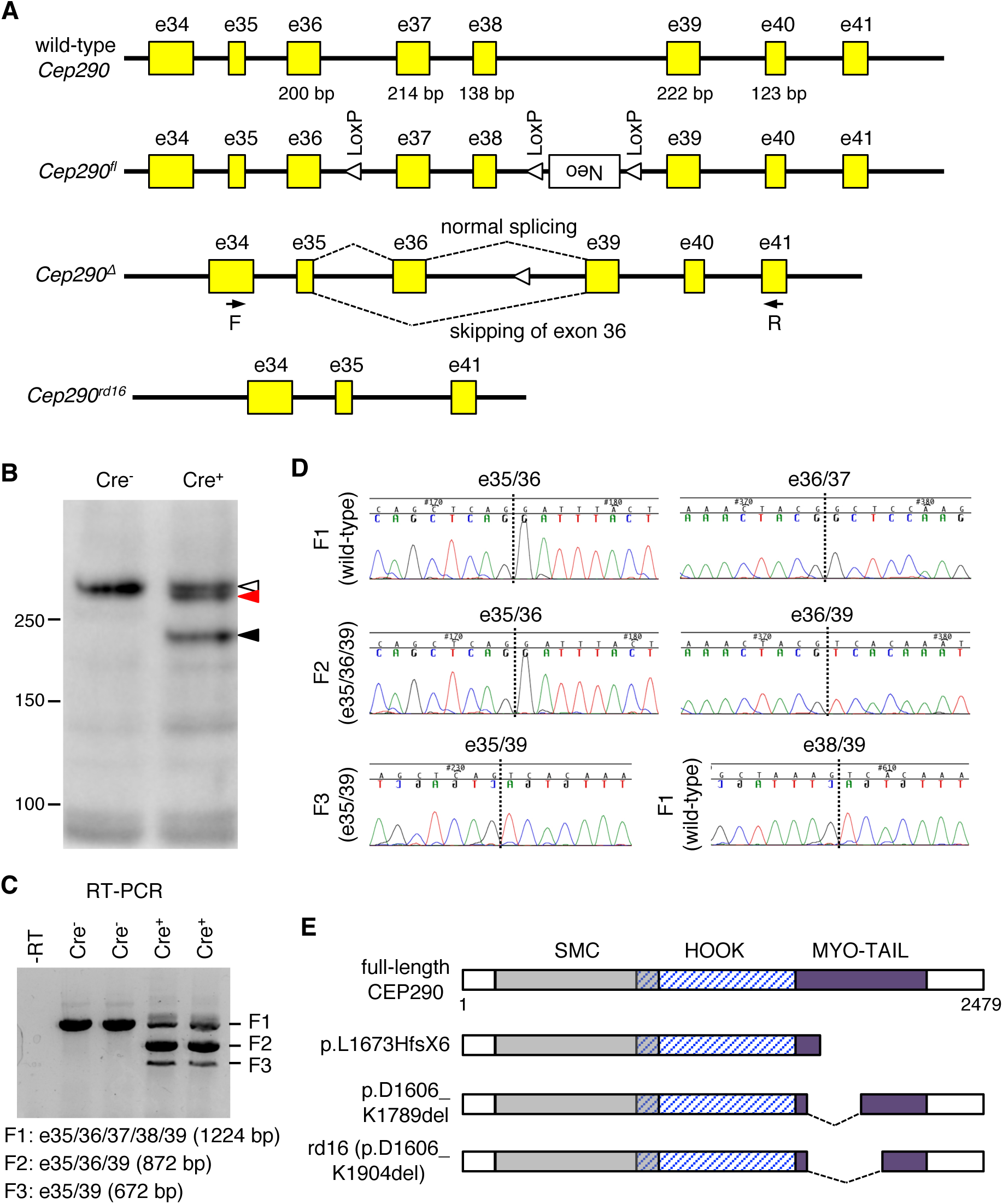
*Cep290*^*Δ*^ is a hypomorphic allele encoding two species of CEP290 deletion mutants. (A) Schematics of *Cep290* allele structures. Yellow boxes represent *Cep290* exons. Exon numbers are described above the yellow boxes. Numbers below yellow boxes indicate the sizes of respective exons (in base pair (bp)). Locations of LoxP sites (open triangles) and a Neo cassette (white box) are marked. Dotted lines depict splicing events between exons 35, 36, and 39 in the *Cep290*^*Δ*^ allele. Black arrows indicate the binding sites of the primers used for RT-PCR (F: forward; R: reverse). (B) Two mutant forms (black and red arrowheads) of CEP290 are detected by CEP290 immunoblotting in *Cep290*^*fl/fl*^*;Cre*^*+*^ retinas. *Cep290*^*fl/fl*^*;Cre*^*-*^ littermates were used as a wild-type control. Open arrowhead indicates full-length CEP290. (C) The *Cep290*^*Δ*^ allele produces two species of mRNAs. Predicted sizes of RT-PCR products (F1, F2, and F3) are shown. A no-reverse-transcriptase control (-RT) was used as a negative control. Each lane represents individual animals. (D) Chromatogram of RT-PCR product sequencing reactions around exon-exon junctions. Vertical dotted lines mark exon-exon junctions. (E) Schematic representation of CEP290 protein products encoded by wild-type, *Cep290*^*Δ*^, and *Cep290*^*rd16*^ alleles. Major structural domains are depicted (SMC: structural maintenance of chromosomes protein homology domain; MYO-TAIL: myosin-tail homology domain). The *Cep290*^*Δ*^ allele produces two species of CEP290 deletion mutants, p.L1673HfsX6 and p.D1606_K1789del.

Since the protein product from the *Cep290*^*Δ*^ allele has not been determined, we extracted proteins from *Cep290*^*fl/fl*^*;Cre*^*-*^ (i.e. control) and *Cep290*^*fl/fl*^*;Cre*^*+*^ retinas at post-natal day (P) 21 and performed CEP290 immunoblotting (**Figure 1B**). Immunoblotting results revealed the production of ∼210 kDa proteins in *Cep290*^*fl/fl*^*;Cre*^*+*^ retinas, which is close to the predicted 196 kDa C-terminally truncated protein (**Figure 1B**; black arrowhead). The presence of full-length CEP290 (open arrowhead) in *Cep290*^*fl/fl*^*;Cre*^*+*^ retinas is expected, because *Cre* is expressed only in rods and other cells in the retina (including cones) produce full-length CEP290. Interestingly, we noticed that there was an additional protein band (red arrowhead) slightly smaller than full-length CEP290. Basal exon skipping and nonsense-associated altered splicing have been observed in human *CEP290* [28–31]. By these mechanisms, exons containing nonsense or frameshift mutations are skipped and the resulting mutant mRNAs maintain the reading frame to produce near-full-length CEP290 proteins instead of being degraded or producing severely truncated proteins. In mouse *Cep290*, we noticed that omitting exon 36 in the *Cep290*^*Δ*^ allele prevents frameshift (**Figure 1A**). To test whether basal exon skipping or nonsense-associated altered splicing occurs, we analyzed *Cep290* mRNAs from *Cep290*^*fl/fl*^*;Cre*^*-*^ and *Cep290*^*fl/fl*^*;Cre*^*+*^ retinas. Reverse transcription PCR (RT-PCR) with primers specific to exons 34 and 41 (forward and reverse primers, respectively) produced a single fragment in controls but 3 fragments in *Cep290*^*fl/fl*^*;Cre*^*+*^ retinas (**Figure 1C**). Sequence analyses of the PCR products revealed that the largest fragment (F1; 1224 bp) was derived from unexcised alleles and contained all exons between exons 34 and 41 (**Figure 1D**). F2 (872 bp), which was the main PCR product from the *Cep290*^*fl/fl*^*;Cre*^*+*^ retinas, contained exons 35, 36, and 39 but not 37 and 38. This indicates that F2 is derived from the *Cep290*^*Δ*^ allele. F3 (672 bp) was devoid of exon 36 in addition to exons 37 and 38, indicating that F3 was derived from the *Cep290*^*Δ*^ allele but exon 36 was skipped during splicing. Since no smaller PCR product was detected in control retinas, our data suggest that exon 36 skipping is a result of nonsense-associated altered splicing rather than basal exon skipping. Loss of exons 36-38 (i.e. 552 bp) causes an in-frame deletion of 184 amino acids (∼20 kDa), which are part of the region deleted in *rd16* mutants (**Figure 1E**). RT-PCR data also suggest that CRE-mediated excision of exons 37 and 38 occurs before P21 in the vast majority of rods. In addition, it is noteworthy that F2 is at least 12-fold more abundant than F3 but the quantity of protein products derived from these transcripts (black and red arrowheads in **Figure 1B**) are comparable. Therefore, the near-full-length in-frame deletion mutant (p.D1606_K1789del) appears to be significantly more stable than the truncated mutant (p.L1673HfsX6). In summary, our data show that *Cep290*^*Δ*^ is a hypomorphic allele encoding two species of mutant proteins, in which the myosin-tail homology domain is disrupted.

### Localization of outer segment-resident proteins in *Cep290*^*fl/fl*^*;Cre*^*+*^ retinas

The myosin-tail homology domain of CEP290 was previously shown to have a microtubule-binding activity [32] and interact with Raf-1 kinase inhibitory protein (RKIP) [33]. *CEP290*-associated ciliopathy patients with mutations in this domain present with more severe phenotypes than predicted by a model based on the total quantity of full-length or near-full-length CEP290 proteins [29]. In addition, disruption of this domain in *rd16* mice causes outer segment biogenesis defects and rapid degeneration of photoreceptors [14]. These findings suggest that the myosin-tail homology domain is essential for CEP290’s function and possibly involved in the trafficking and confinement of outer segment proteins. Therefore, we examined localization of outer segment-resident proteins in *Cep290*^*fl/fl*^*;Cre*^*+*^ conditional mutant mice.

To this end, we probed localization of RHO, PRPH2, ROM1, ABCA4, PDE6B, GUCY2D, ATP8A2, and CNGA1 in *Cep290*^*+/fl*^*;Cre*^*+*^ (control) and *Cep290*^*fl/fl*^*;Cre*^*+*^ retinas. Whilst RHO localizes to both disc membranes and outer segment plasma membranes [34, 35], PRPH2, ROM1, ABCA4, GUCY2D, and ATP8A2 specifically localize to disc membranes [36–41]. CNGA1 specifically localizes to the outer segment plasma membrane [42], which is contiguous with the inner segment plasma membrane (hereafter our use of the term inner segment encompasses all parts of photoreceptors including ellipsoid, myoid, soma, axon, and synaptic terminal but excluding the outer segment). At P11, when the connecting cilium assembly is completed and the outer segment is rapidly growing [43], no significant differences were observed between control and *Cep290*^*fl/fl*^*;Cre*^*+*^ retinas with respect to the localization of outer segment-resident proteins, including RHO (**Supplementary Figure S1**). At P20, most outer segment-resident proteins examined still showed normal localization to the outer segment (**Figure 2**). RHO was the only exception and significant mislocalization was observed in inner segments (**Figure 2A**), indicating that RHO mislocalization is one of the earliest events that occur upon disruption of the myosin-tail homology domain.

**Figure 2.**
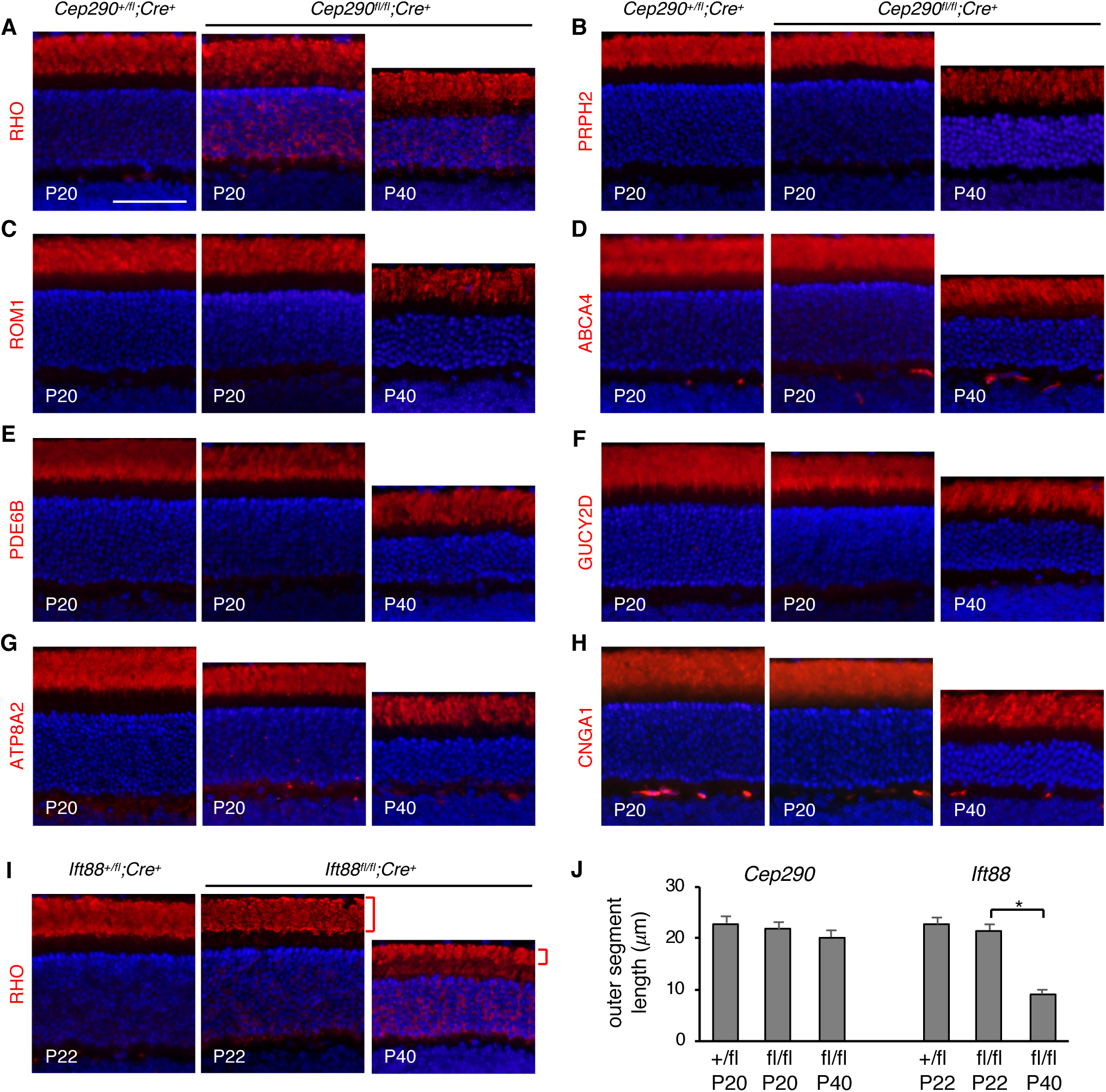
Localization of outer segment-resident proteins in *Cep290*^*fl/fl*^*;Cre*^*+*^ mice. Localization of RHO (A), PRPH2 (B), ROM1 (C), ABCA4 (D), PDE6B (E), GUCY2D (F), ATP8A2 (G), and CNGA1 (H) was probed in *Cep290*^*+/fl*^*;Cre*^*+*^ (at P20) and *Cep290*^*fl/fl*^*;Cre*^*+*^ (at P20 and P40) retinas. (I) Retinal sections of *Ift88*^*+/fl*^*;Cre*^*+*^ (at P22) and *Ift88*^*fl/fl*^*;Cre*^*+*^ (at P22 and P40) mice were immuno-stained with anti-RHO antibody and shown for comparisons with *Cep290*^*fl/fl*^*;Cre*^*+*^ mice. Red brackets indicate outer segments. Sections were counterstained with 4′,6-Diamidine-2′-phenylindole dihydrochloride (DAPI) to show nuclei (blue). At least 3 animals, both male and female, were used per group and representative images are shown. Scale bar represents 50 μm. (J) Length of the outer segment in *Cep290*^*fl/fl*^*;Cre*^*+*^ and *Ift88*^*fl/fl*^*;Cre*^*+*^ retinas. Lengths of the outer segment were measured in areas 0.5-1.0 mm away from the optic nerve. Data are from 3 animals per genotype (2 sections per animal). Mean and standard deviation (SD; error bar) are shown. Asterisk indicates statistical significance (two-tailed, two-sample *t*-test assuming unequal variances; *p*<0.01).

To test whether outer segment protein localization deteriorates as photoreceptors degenerate, we examined the localization of the above 8 proteins at P40 (**Figure 2**). Twenty days (from P20 to P40) is sufficient for the outer segment to renew entirely in mouse retinas [2], and approximately 30-40% of photoreceptors were lost by P40 in *Cep290*^*fl/fl*^*;Cre*^*+*^ retinas. However, no significant mislocalization of PRPH2, ROM1, ABCA4, PDE6B, GUCY2D, ATP8A2, and CNGA1 was observed. In addition, the length of the outer segment was not significantly reduced despite the progression of degeneration and cell loss. This observation sharply contrasts with the rapid shortening of the outer segment in *Ift88*^*fl/fl*^*;Cre*^*+*^ mice, in which intraflagellar transport (IFT) to maintain the outer segment is ablated, between P22 (when RHO mislocalization is first noticeable) and P40 (red bracket; **Figure 2I and J**). Therefore, most outer segment proteins except RHO appear to be properly transported and confined to the outer segment in *Cep290*^*fl/fl*^*;Cre*^*+*^ rods.

### Accumulation of inner segment membrane proteins in the outer segment upon disruption of the CEP290 myosin-tail homology domain

We then examined whether disruption of the myosin-tail homology domain affected confinement of inner segment membrane proteins. STX3 is a SNARE (<u>S</u>oluble <u>N</u>-ethymaleimide-sensitive factor <u>A</u>ttachment protein <u>RE</u>ceptor) protein with a single transmembrane domain at its C-terminus and facilitates membrane/vesicle fusion for exocytosis. In normal photoreceptors, STX3 is found in the plasma membrane throughout the inner segment but not in the outer segment [44, 45] (**Figure 3A**; left). Since STX3 and its interacting partner STXBP1 mislocalize to the outer segment in Bardet-Biedl syndrome (BBS) mutant retinas [45–47], we speculated that these proteins might need an intact ciliary gate for their inner segment-restricted localization. Indeed, STX3 and STXBP1 were found mislocalized to outer segments in *Cep290*^*fl/fl*^*;Cre*^*+*^ retinas at P20 (**Figure 3A and B**; also see **Figure 3E** for quantification).

**Figure 3.**
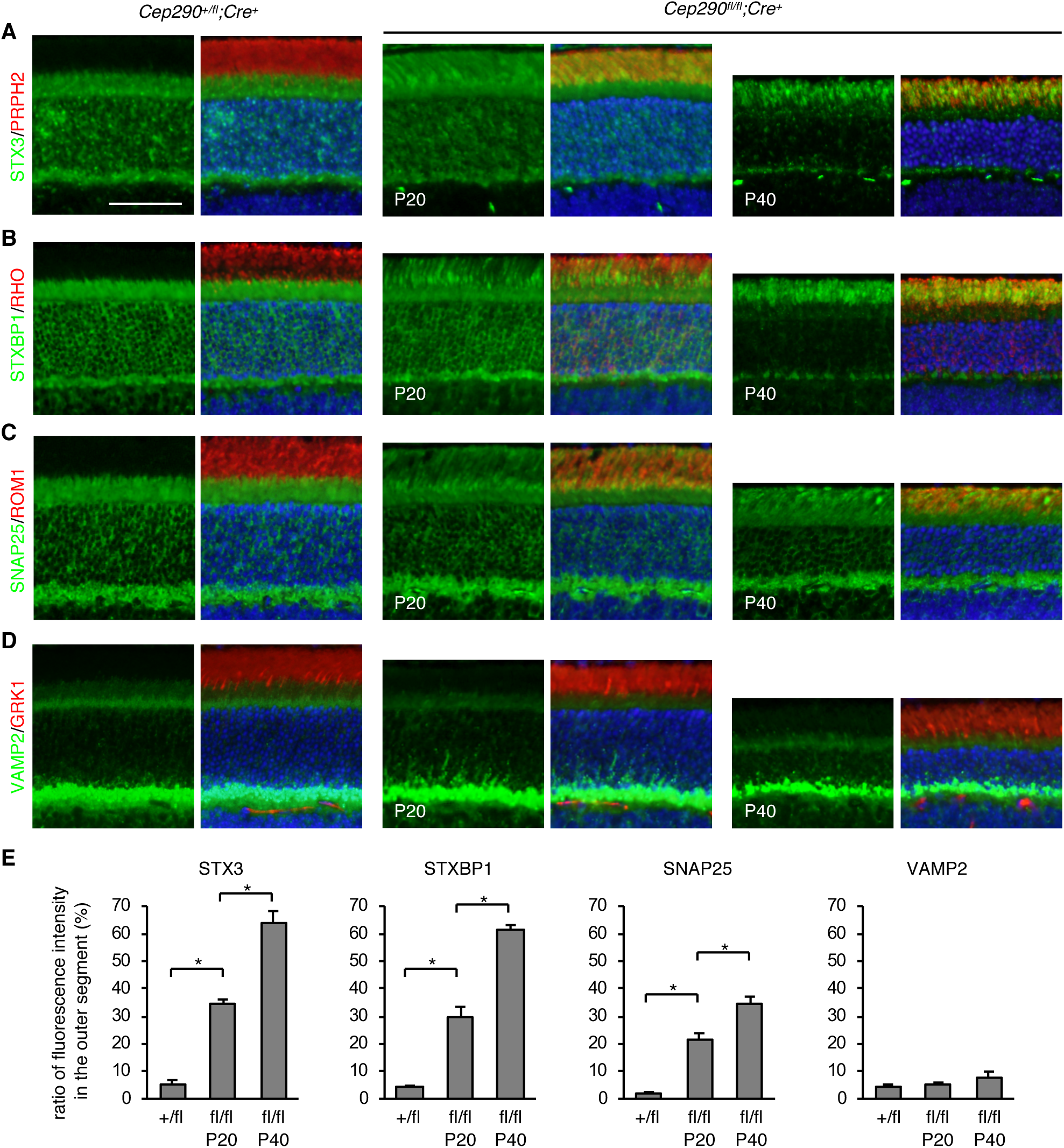
Localization of STX3 and its interacting proteins in *Cep290*^*fl/fl*^*;Cre*^*+*^ mice. Control (*Cep290*^*+/fl*^*;Cre*^*+*^: at P20) and *Cep290*^*fl/fl*^*;Cre*^*+*^ (at P20 and P40) mouse retinal sections were immuno-stained with STX3 (A), STXBP1 (B), SNAP25 (C), and VAMP2 (D) antibodies (green). To delineate outer segments, sections were co-stained with PRPH2, RHO, ROM1, and GRK1 antibodies (red). DAPI was used to label nuclei (blue). Merged images are shown on the right. At least 3 animals, both male and female, were used per group and representative images are shown. Scale bar represents 50 μm. (E) Quantification of inner segment protein mislocalization to the outer segment. Depicted are ratios of integrated fluorescence intensities in the outer segment relative to the photoreceptor cell layer. Data are from 3 mice (2 sections per mouse and 2 areas per section). Mean and standard deviation (SD; error bar) are shown. Asterisks indicate statistical significance (two-tailed, two-sample *t*-test assuming unequal variances; *p*<0.01).

These results prompted us to examine the localization of other inner segment-specific membrane proteins. SNAP25 and VAMP2 are SNARE proteins that interact with STX3. Although both SNAP25 and VAMP2 are restricted to the inner segment, SNAP25 is a t-SNARE protein localizing to the plasma membrane while VAMP2 is a v-SNARE protein present on secretory vesicles. Also, while SNAP25 is almost evenly distributed throughout the inner segment, VAMP2 is highly enriched at synaptic terminals (**Figure 3C and D**; left). In 20-day old *Cep290*^*fl/fl*^*;Cre*^*+*^ mice, SNAP25 was found dispersed throughout the cell including the outer segment (**Figure 3C and E**). In contrast, VAMP2 localization was not altered in *Cep290*^*fl/fl*^*;Cre*^*+*^ rods (**Figure 3D and E**).

IMPG2 (interphotoreceptor matrix proteoglycan 2) is a single transmembrane domain-containing proteoglycan found in the interphotoreceptor matrix, an extracellular matrix between the retinal pigmented epithelium (RPE) and the outer limiting membrane of the retina [48, 49]. Mutations in IMPG2 are associated with retinitis pigmentosa and vitelliform macular dystrophy [50–52]. We previously identified IMPG2 as one of the proteins enriched in *Lztfl1* mutant outer segments [45]. When probed with an antibody raised to its C-terminal intracellular domain, IMPG2 immunoreactivity was mostly found within the ellipsoid and myoid zones of the inner segment in normal photoreceptors (**Figure 4A**; also see **Supplementary Figure S2** for antibody characterization and discussion). In *Cep290*^*fl/fl*^*;Cre*^*+*^ retinas, however, a significant amount of IMPG2 immunoreactivity was detected in outer segments in addition to inner segments (**Figure 4A and F**).

**Figure 4.**
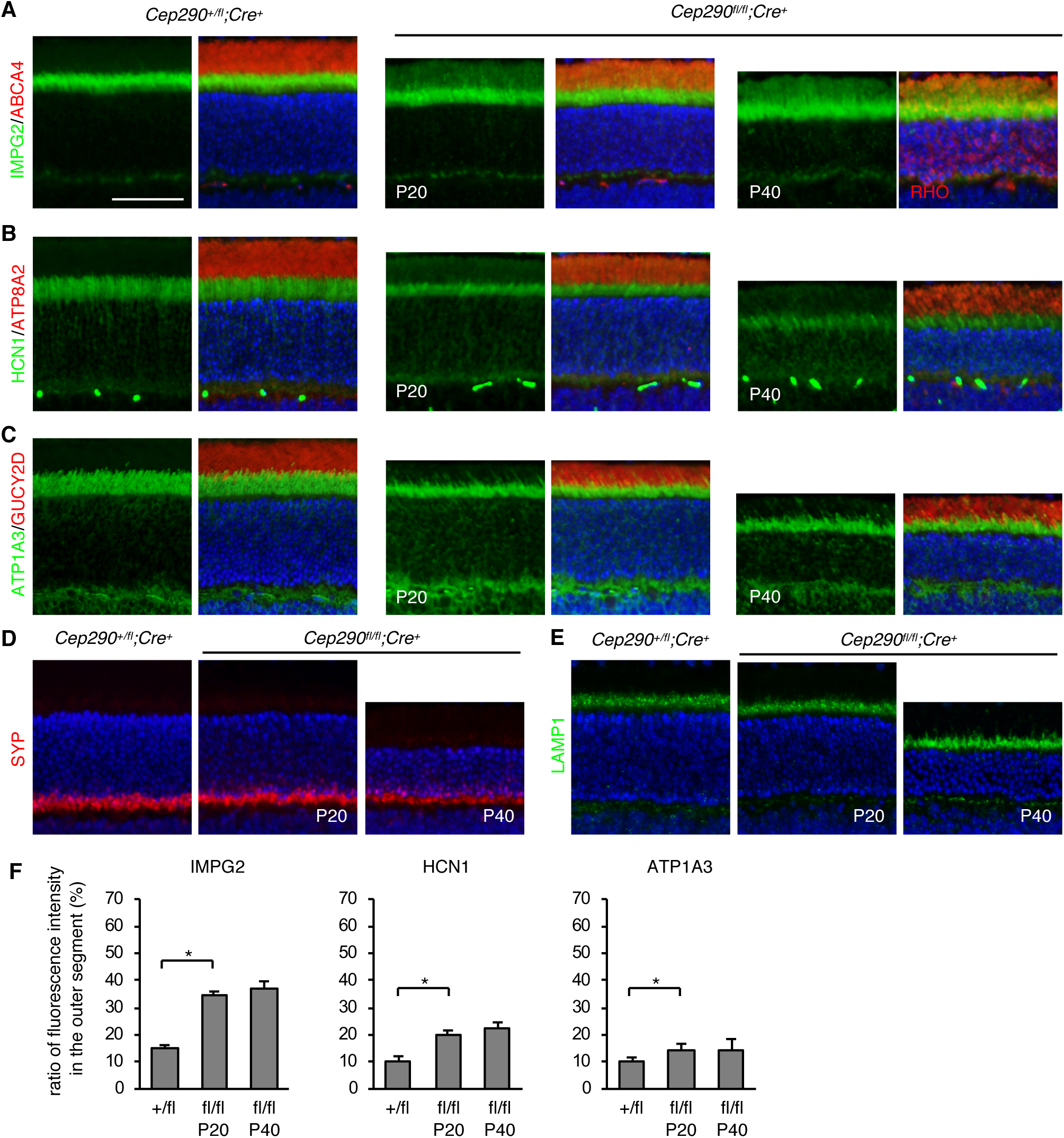
Localization of inner segment membrane proteins in *Cep290*^*fl/fl*^*;Cre*^*+*^ mice. Retinal sections from *Cep290*^*+/fl*^*;Cre*^*+*^ (at P20) and *Cep290*^*fl/fl*^*;Cre*^*+*^ (at P20 and P40) mice were labelled with IMPG2 (A), HCN1 (B), and ATP1A3 (C) antibodies (green). Outer segments were delineated with ABCA4 or RHO (A), ATP8A2 (B), and GUCY2D (C) antibodies (red). Nuclei were counterstained with DAPI. (D-E) Localization of endomembrane proteins SYP (D) and LAMP1 (E) was examined using respective antibodies. At least 3 animals were used per group and representative images are shown. Scale bar represents 50 μm. (F) Quantification of inner segment membrane protein mislocalization to the outer segment. Others are the same as in Figure 3.

HCN1 (K^+^/Na^+^ hyperpolarization-activated cyclic nucleotide-gated channel 1) and ATP1A3 (Na^+^/K^+^-transporting ATPase1 subunit *α*3) are transmembrane proteins localizing to the inner segment plasma membrane [53–55]. Within the inner segment, HCN1 and ATP1A3 are enriched within the myoid and ellipsoid zones (**Figure 4B, C, and F**). Mild mislocalization of HCN1 was observed throughout the outer segment in *Cep290*^*fl/fl*^*;Cre*^*+*^ retinas. ATP1A3 also exhibited mild mislocalization to outer segments but there was a considerable cell-to-cell variation with respect to the severity of mislocalization. SYP (synaptophysin) and LAMP1 (lysosome-associated membrane glycoprotein 1) are integral membrane proteins that specifically localize to synaptic vesicles and lysosomes, respectively. As expected, SYP was found at synaptic terminals and LAMP1 was in the ellipsoid zone in normal photoreceptors. Localization of these endomembrane proteins was not altered in *Cep290*^*fl/fl*^*;Cre*^*+*^ retinas (**Figure 4D and E**).

To test whether mislocalization of inner segment proteins becomes more prevalent or severe as retinal degeneration progresses, we examined the localization of the aforementioned proteins at P40 (**Figures 3 and 4**; right panels). By this age, the majority (∼60%) of STX3 and STXBP1 localized to the outer segment (see **Figure 3E** for quantification). It should be noted that this is an underestimation, because *Stx3* and *Stxbp1* are expressed in not only rods but also cones and residual signals in the synaptic terminals (which are included in quantification as a part of the inner segment) are mostly from cones, in which *Cre* is not expressed (see **Supplementary Figure S3)**. SNAP25 showed a moderate increase in mislocalization at P40, but the signal intensity in the outer segment did not exceed that in the inner segment. Mislocalization of other inner segment proteins was not appreciably exacerbated from what was observed at P20. These data show that the myosin-tail homology domain of CEP290 is essential for confinement of inner segment membrane proteins. Our data also suggest that proteins on the plasma membranes are primarily subject to diffusion and that STX3 and STXBP1 are particularly susceptible to accumulation in the outer segment.

### Mislocalization of inner segment membrane proteins in *Cep290*^*fl/fl*^*;Cre*^*+*^ retinas is not due to loss of BBSome functions

Accumulation of STX3 and STXBP1 in the outer segment is a phenotype commonly observed in BBS mutant photoreceptors [45–47]. While CEP290 is a stationary protein constituting the ciliary gate, BBSomes (a protein complex composed of 8 BBS proteins [56, 57]) are adapter complexes that link ciliary cargos to IFT particles and transport them in and out of the ciliary compartment [58–65]. To test whether STX3 and STXBP1 mislocalization in *Cep290*^*fl/fl*^*;Cre*^*+*^ retinas is mediated by loss or dysfunction of the BBSome, we examined the quantity and localization of BBSome components as well as a BBSome regulator LZTFL1 [66]. Immunoblot analyses of retinal extracts from 25-day-old *Cep290*^*+/fl*^*;Cre*^*?*^ (control) and *Cep290*^*fl/fl*^*;Cre*^*+*^ mice indicated that there was a slight to moderate (12-25%) reduction of photoreceptor-specific proteins (RHO, GRK1, PRPH2, and PDE6A) in *Cep290*^*fl/fl*^*;Cre*^*+*^ retinas (**Figure 5A**). This is consistent with some loss of photoreceptors by this age. BBSome components (BBS2 and BBS7) and LZTFL1 showed similar levels of reduction. These data suggest that reduction of BBS proteins is proportionate to the loss of photoreceptors and that BBS protein levels are not affected by impaired CEP290 functions in *Cep290*^*fl/fl*^*;Cre*^*+*^ retinas. We then examined BBSome assembly by immunoprecipitation. As shown in **Figure 5B**, all BBSome components tested were similarly pulled down by BBS7 antibodies in control and *Cep290*^*fl/fl*^*;Cre*^*+*^ retinas. These data indicate that BBSome assembly is not altered in *Cep290*^*fl/fl*^*;Cre*^*+*^ retinas.

**Figure 5.**
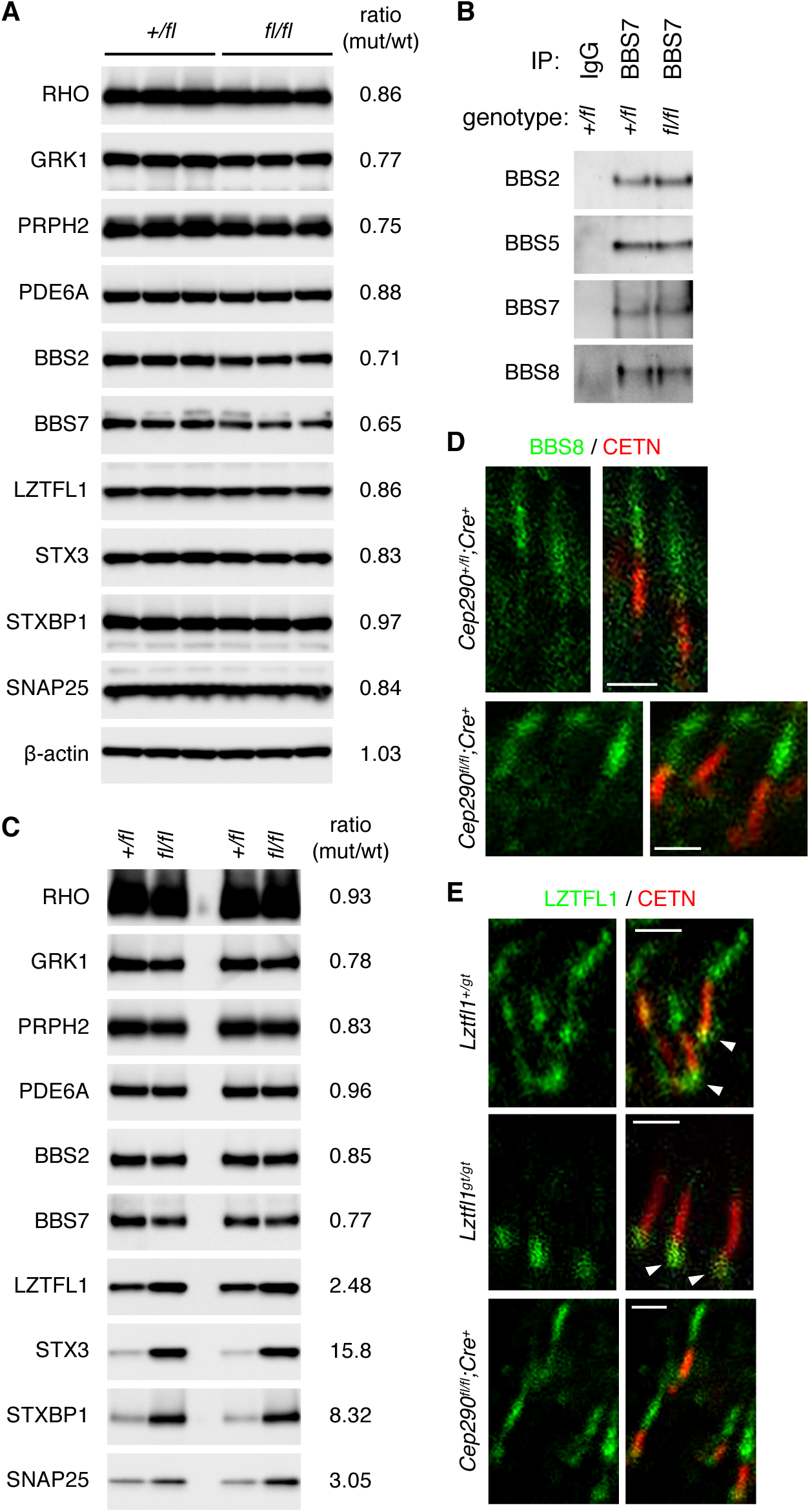
Expression and localization of BBS proteins is not altered in *Cep290*^*fl/fl*^*;Cre*^*+*^ retinas. (A) Immunoblot analysis of *Cep290*^*+/fl*^*;Cre*^*?*^ and *Cep290*^*fl/fl*^*;Cre*^*+*^ retinal protein extracts. Each lane represents individual animals and 3 animals were analyzed per genotype. Thirty μg of proteins were loaded per lane. Numbers on the right are relative band intensities of the indicated proteins in *Cep290*^*fl/fl*^*;Cre*^*+*^ retinas compared to those in *Cep290*^*+/fl*^*;Cre*^*?*^ retinas. (B) BBSome assembly is not altered in *Cep290*^*fl/fl*^*;Cre*^*+*^ retinas. Retinal protein extracts from *Cep290*^*+/fl*^*;Cre*^*?*^ and *Cep290*^*fl/fl*^*;Cre*^*+*^ retinas were subjected to immunoprecipitation (IP) with anti-BBS7 antibody. Normal rabbit IgG was used as a negative control. Co-precipitated BBS proteins were detected by immunoblotting. (C) Immunoblot analysis of outer segment fractions isolated from *Cep290*^*+/fl*^*;Cre*^*?*^ and *Cep290*^*fl/fl*^*;Cre*^*+*^ retinas. Each lane represents individual outer segment preparations and results from two independent preparations are shown. Five μg of proteins were loaded per lane. Others are same as in (A). (D) Localization of BBS8 (green) in *Cep290*^*+/fl*^*;Cre*^*+*^ and *Cep290*^*fl/fl*^*;Cre*^*+*^ photoreceptors. Connecting cilia were labeled with anti-CETN antibody (red). Scale bar represents 1 μm. (E) Localization of LZTFL1 (green) in control (*Lztfl1*^*+lgt*^), *Lztfl1*^*gt/gt*^, and *Cep290*^*fl/fl*^*;Cre*^*+*^ photoreceptors. White arrowheads indicate binding of anti-LZTFL1 antibody to cross-reacting protein(s) around the basal body. Others are same as in (D).

To assess the quantity of BBS proteins within the outer segment, we isolated outer segments from *Cep290*^*+/fl*^*;Cre*^*?*^ and *Cep290*^*fl/fl*^*;Cre*^*+*^ retinas and conducted immunoblot analyses (**Figure 5C**). After normalization to total protein quantities, no significant differences were observed between normal and CEP290-deficient outer segments with respect to the quantity of outer segment-resident proteins. BBSome components, BBS2 and BBS7, also showed no significant differences. Interestingly, however, there was a more than 2-fold increase in LZTFL1 quantity in outer segments from *Cep290*^*fl/fl*^*;Cre*^*+*^ retinas. Consistent with the mislocalization of STX3 and STXBP1 in the outer segment, there was a large increase of these proteins in *Cep290*^*fl/fl*^*;Cre*^*+*^ outer segments. SNAP25 showed ∼3 fold increase in *Cep290*^*fl/fl*^*;Cre*^*+*^ outer segments.

We then examined the localization of BBS proteins in normal and *Cep290*^*fl/fl*^*;Cre*^*+*^ photoreceptors. While CEP290 is localized to the connecting cilium, BBS5 was previously shown to localize along the axoneme in the outer segment [67]. BBS5 was also detected within the connecting cilium at a lower intensity. Very similar localization patterns were observed with BBS8 and LZTFL1 (**Figure 5D and E**). LZTFL1 immunoreactivity was also detected around the basal body but this staining persisted in *Lztfl1* mutant [45] retinas, indicating that signals around the basal bodies are from cross-reacting protein(s) (**Figure 5E;** arrowheads). Localization of BBS8 and LZTFL1 in *Cep290*^*fl/fl*^*;Cre*^*+*^ retinas was comparable to that of normal photoreceptors (**Figure 5D and E**). These data indicate that, despite some quantitative changes in LZTFL1 in the outer segment, overall localization patterns of the BBSome and LZTFL1 are not altered in *Cep290*^*fl/fl*^*;Cre*^*+*^ retinas. Therefore, we conclude that the accumulation of STX3 and STXBP1 in the outer segment in *Cep290*^*fl/fl*^*;Cre*^*+*^ retinas is unlikely due to alterations in BBSome functions.

### Accumulation of inner segment membrane proteins in the outer segment in *rd16* mice

We next examined whether similar protein mislocalization was observed in *rd16* mice (**Figure 6**). As previously described [14], retinal degeneration in *rd16* mice was already evident at P15 and outer segments were rudimentary, suggesting that outer segment biogenesis is severely impaired in *rd16* mice. RHO mislocalization was also evident in the inner segment (**Figure 6B**). In addition, presumably due to the outer segment biogenesis defect, low level mislocalization of outer segment proteins was detected in general. However, the highest immunoreactivity of all outer segment-resident proteins examined was detected in the outer segment, suggesting that these proteins are still delivered to the outer segment in *rd16* mice. Mislocalization of CNGA1 was relatively obvious compared to other outer segment proteins and predominantly restricted to the distal (ellipsoid) portion of the inner segment as opposed to being dispersed throughout the inner segment like RHO. (**Figure 6H**).

**Figure 6.**
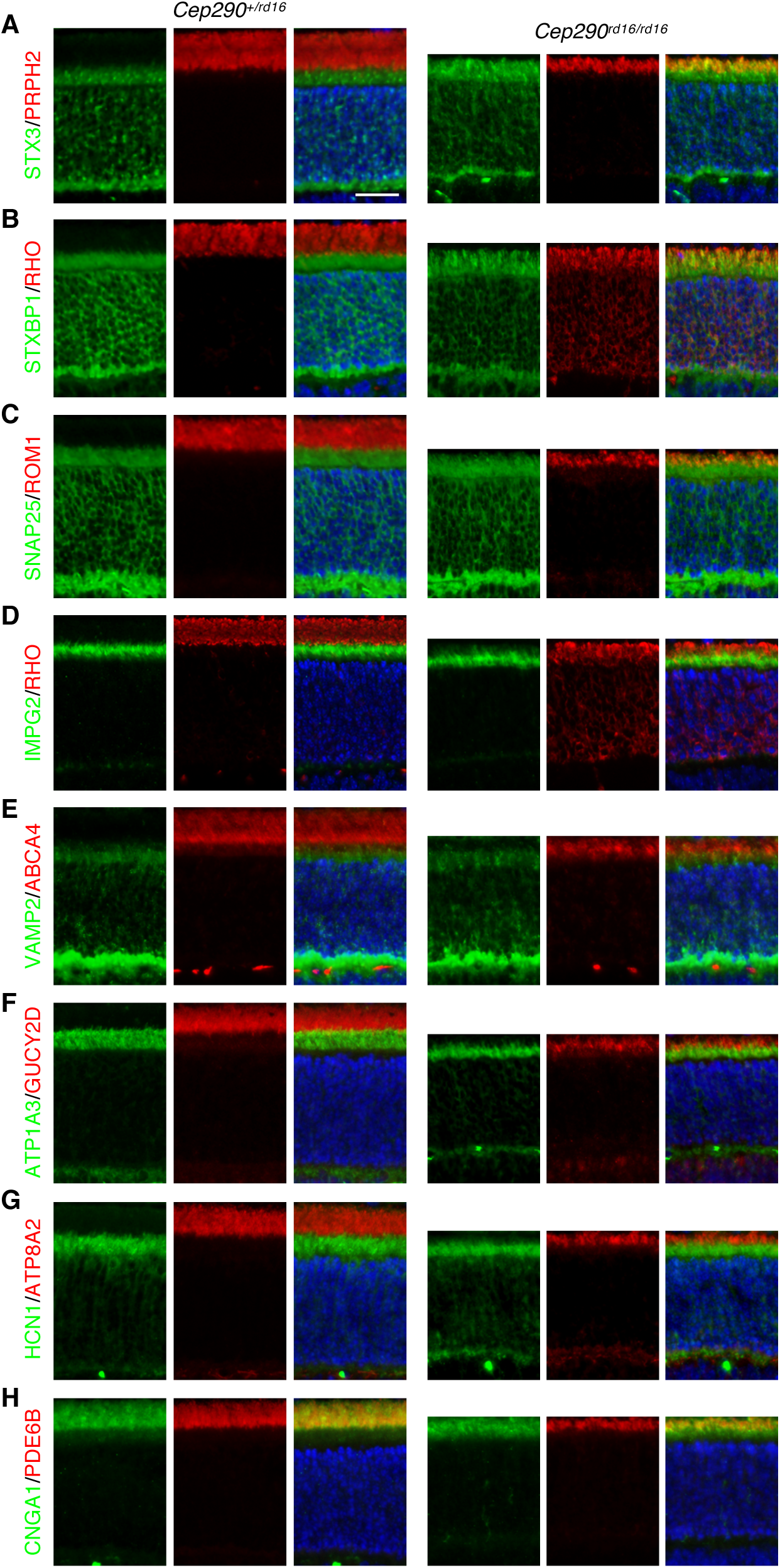
Disruption of compartmentalized protein localization in *Cep290*^*rd16/rd16*^ mice. Localization of outer segment-resident proteins (PRPH2, RHO, ROM1, ABCA4, GUCY2D, ATP8A2, PDE6B, and CNGA1; all in red except CNGA1, which is in green) and inner segment membrane proteins (STX3, STXBP1, SNAP25, IMPG2, VAMP2, ATP1A3, and HCN1; all in green) was examined by indirect immunofluorescence microscopy. Scale bar represents 25 μm.

We then examined the localization of inner segment membrane proteins in *rd16* mice. Proteins that exhibited significant mislocalization in *Cep290*^*fl/fl*^*;Cre*^*+*^ retinas (STX3, STXBP1, SNAP25, and IMPG2) all showed obvious mislocalization to the outer segment in *rd16* retinas (**Figure 6**). In contrast, localization of VAMP2, ATP1A3, HCN1, SYP, and LAMP1 was not noticeably altered in *rd16* retinas (**Figure 6** and data not shown). These results indicate that protein mislocalization phenotypes are comparable in *Cep290*^*fl/fl*^*;Cre*^*+*^ and *rd16* retinas and that similar pathomechanisms underlie retinal degeneration in *rd16* mice as in *Cep290*^*fl/fl*^*;Cre*^*+*^ mice.

## DISCUSSION

Mutations that cause *CEP290*-associated ciliopathies are found throughout the gene. The vast majority of them are truncating mutations (i.e. nonsense or frameshift mutations) [22]. This means that most mutant alleles are destined to produce either no proteins (if mutant mRNAs are degraded by nonsense-mediated decay) or more likely C-terminally truncated proteins. Certain truncating mutations avoid these detrimental consequences by skipping exons and producing near-full-length proteins, resulting in unexpectedly mild phenotypes [28–31]. The myosin-tail homology domain is within the C-terminal 1/3 of CEP290, and therefore not produced in a large proportion of *CEP290* mutant alleles with truncating mutations. The mouse models used in our study represent *CEP290*-associated retinopathies in which the myosin-tail homology domain is disrupted. *rd16* mice have an in-frame deletion of 299 amino acids (aa 1606-1904) within this domain. Rods in *Cep290*^*fl/fl*^*;Cre*^*+*^ mice are effectively compound heterozygotes expressing two deletion mutants, p.L1673HfsX6 and p.D1606_K1789del. Our study demonstrates that CEP290, more specifically the myosin-tail homology domain of CEP290, is essential for protein confinement between the inner and the outer segments and thus maintains compartmentalized protein localization in photoreceptors. Disruption of this domain and consequent deficits in CEP290 function cause encroachment of select inner segment plasma membrane proteins on the outer segment and mislocalization of RHO to the inner segment. Therefore, our study supports the current model of CEP290 function as a ciliary gatekeeper and identifies disruption of compartmentalized protein localization as part of the disease mechanisms underlying *CEP290*-associated retinopathies.

Our study shows that the myosin-tail homology domain is essential for CEP290’s function as a ciliary gatekeeper, particularly to prevent diffusion of inner segment membrane proteins into the outer segment. A subset of inner segment membrane proteins rapidly accumulate in the outer segment upon disruption of the myosin-tail homology domain. STX3 and STXBP1, in particular, show striking accumulation in outer segments as degeneration progresses. Other inner segment membrane proteins tested show various degrees of accumulation in the outer segment. SNAP25 and IMPG2 exhibit significant accumulation in the outer segment but their density in the outer segment does not exceed that in the inner segment. It is tantalizing to speculate that there are additional factor(s) that induce STX3 and STXBP1 enrichment in the outer segment. Mislocalization of ATP1A3 and HCN1 is mild, and localization of VAMP2, LAMP1, and SYP is not affected by the loss of CEP290 myosin-tail homology domain.

In contrast, the myosin-tail homology domain appears to be dispensable for the trafficking and confinement of most outer segment-resident proteins except RHO. In *rd16* mice, the myosin-tail homology domain is constitutively disrupted and the outer segment biogenesis is severely impaired. Most outer segment proteins, however, manage to be delivered to the rudimentary outer segment. The low level mislocalization of outer segment proteins is likely secondary to the outer segment biogenesis defects including small size and disorganized disc structures. In *Cep290*^*fl/fl*^*;Cre*^*+*^ retinas, in which the myosin-tail homology domain is disrupted after the connecting cilium is formed, mislocalization of outer segment-resident proteins is not observed except for RHO. Furthermore, the length of the outer segment does not change significantly for 20 days while degeneration is progressing. Zebrafish *cep290*^*fh297/fh297*^ mutants, which have a nonsense mutation (p.Q1217*) before the myosin-tail homology domain, exhibit partial mislocalization of RHO but not of rhodopsin kinase GRK1 or a tranducin subunit GNB1 at 6 months post fertilization [24]. Although degradation of mis-trafficked proteins in the inner segment could contribute to the apparent lack of mislocalized proteins, these data suggest that most outer segment proteins do not require the myosin-tail homology domain for their outer segment-specific localization. It remains to be determined whether the residual part of CEP290, which is expressed in *Cep290*^*fl/fl*^*;Cre*^*+*^ and *rd16* retinas, is sufficient for the trafficking and confinement of outer segment proteins and, if so, specifically which proteins require CEP290 for their outer segment localization. Among the outer segment proteins examined, RHO was the only protein that showed consistent mislocalization in both *Cep290*^*fl/fl*^*;Cre*^*+*^ and *rd16* retinas. It is currently unclear whether mislocalized RHOs originate from the outer segment (i.e. by leakage), whether they are newly synthesized proteins that failed to be delivered, or both. More sophisticated approaches such as a pulse-chase experiment are needed to address this question.

Although the number of proteins examined in the present study is limited, it is noteworthy that all proteins that mislocalize upon disruption of the myosin-tail homology domain are plasma membrane-localizing proteins. For instance, STX3 and SNAP25 are transmembrane proteins known to localize to the inner segment plasma membrane [44, 53–55]. STXBP1 does not have a transmembrane domain or a lipid anchor. However, STXBP1 localization tightly correlates with that of STX3 in both BBS and *Cep290* mutant retinas at all ages tested ([45] and this study). In addition, STXBP1-STX3 protein complexes are readily detectable by immunoprecipitation followed by silver staining in retinal extracts (**Supplementary Figure S4**). These observations suggest that most STXBP1 is likely associated with STX3 in photoreceptors. Therefore, STXBP1 can be considered as a peripheral membrane protein associated with the plasma membrane through STX3. IMPG2 is a single transmembrane protein identified as a constituent of the interphotoreceptor matrix [49]. Since the procedures used in the prior study to isolate IMPG2 from the interphotoreceptor matrix were expected not to extract integral membrane proteins, Acharya *et al*. predicted that IMPG2 might be cleaved before the transmembrane domain to release the N-terminal fragment into the extracellular matrix [49]. Indeed, we found that recombinant IMPG2 expressed in HEK293T cells was efficiently cleaved (**Supplementary Figure S2A**). The C-terminal fragment containing the transmembrane domain and the intracellular domain, to which our anti-IMPG2 antibody was raised, localized to the plasma membrane (**Supplementary Figure S2B**). These data suggest that IMPG2 immunoreactivity detected by our IMPG2 antibody in the retina is from full-length IMPG2 in the secretory pathway (including endoplasmic reticulum and Golgi) or the C-terminal cleavage product that localizes to the plasma membrane. In contrast, localization of endomembrane proteins (VAMP2, SYP, and LAMP1) is not affected by the loss of the myosin-tail homology domain. These proteins are actively sorted and targeted to their destinations upon synthesis and during recycling, and therefore not subject to diffusion along the plasma membrane. ATP1A3 and HCN1 localize to the inner segment plasma membrane but exhibit only mild mislocalization. We speculate that these proteins may be targeted and retained in the inner segment plasma membrane by unknown mechanisms. Indeed, contrary to STX3, STXBP1, and SNAP25, which are evenly distributed throughout the inner segment in normal photoreceptors, ATP1A3 and HCN1 are significantly enriched in the myoid and ellipsoid zones, implying that they do not freely diffuse within the inner segment. Taken together, our data suggest that a subset of inner segment plasma membrane proteins are liable to mislocalization to the outer segment upon disruption of the myosin-tail homology domain.

Based on our study, we propose that failure of protein confinement at the connecting cilium and consequent accumulation of inner segment plasma membrane proteins in the outer segment combined with insufficient RHO delivery underlies retinal degeneration in *CEP290*-associated ciliopathies (**Figure 7**). In normal photoreceptors, CEP290 is a part of the ciliary gate that confines inner segment plasma membrane proteins to the inner segment. The myosin-tail homology domain is crucial for this function. CEP290 is also required for the trafficking and/or confinement of RHO to the outer segment. In photoreceptors with compromised CEP290 functions, ciliary gates are impaired, allowing diffusion of select inner segment plasma membrane proteins into the outer segment. Abundance of membranes in the outer segment is likely to contribute to the accumulation of inner segment membrane proteins. Perhaps, accumulation of inner segment membrane proteins in nascent discs combined with insufficient delivery of RHO disturbs the disc morphogenesis process and causes or contributes to the outer segment biogenesis defects observed in *rd16* mice. Inner segment mislocalization of RHO is likely another key pathomechanism that triggers photoreceptor death, as mutations disrupting RHO trafficking commonly cause retinal degeneration ([68] and references therein). Precise roles of CEP290 for outer segment protein trafficking and confinement as well as the mechanisms by which the myosin-tail homology domain blocks inner segment membrane proteins remain to be determined. It also should be noted that both *CEP290* mouse models used in the present study possess hypomorphic alleles not a null. *Cep290* null mutants completely lack the connecting cilium and the outer segment [15]. Therefore, although not fully understood, one should bear in mind that CEP290 has additional roles in the connecting cilium biogenesis and that the N-terminal 2/3 of CEP290 might be necessary for outer segment protein trafficking and confinement. Finally, ciliary gates are composed of multiple proteins of the “NPHP” and “MKS modules” [5–8, 69]. Loss-of-function mutations in these genes/proteins commonly cause retinal degeneration. Similar pathomechanisms may underlie retinal degenerations in these diseases.

**Figure 7.**
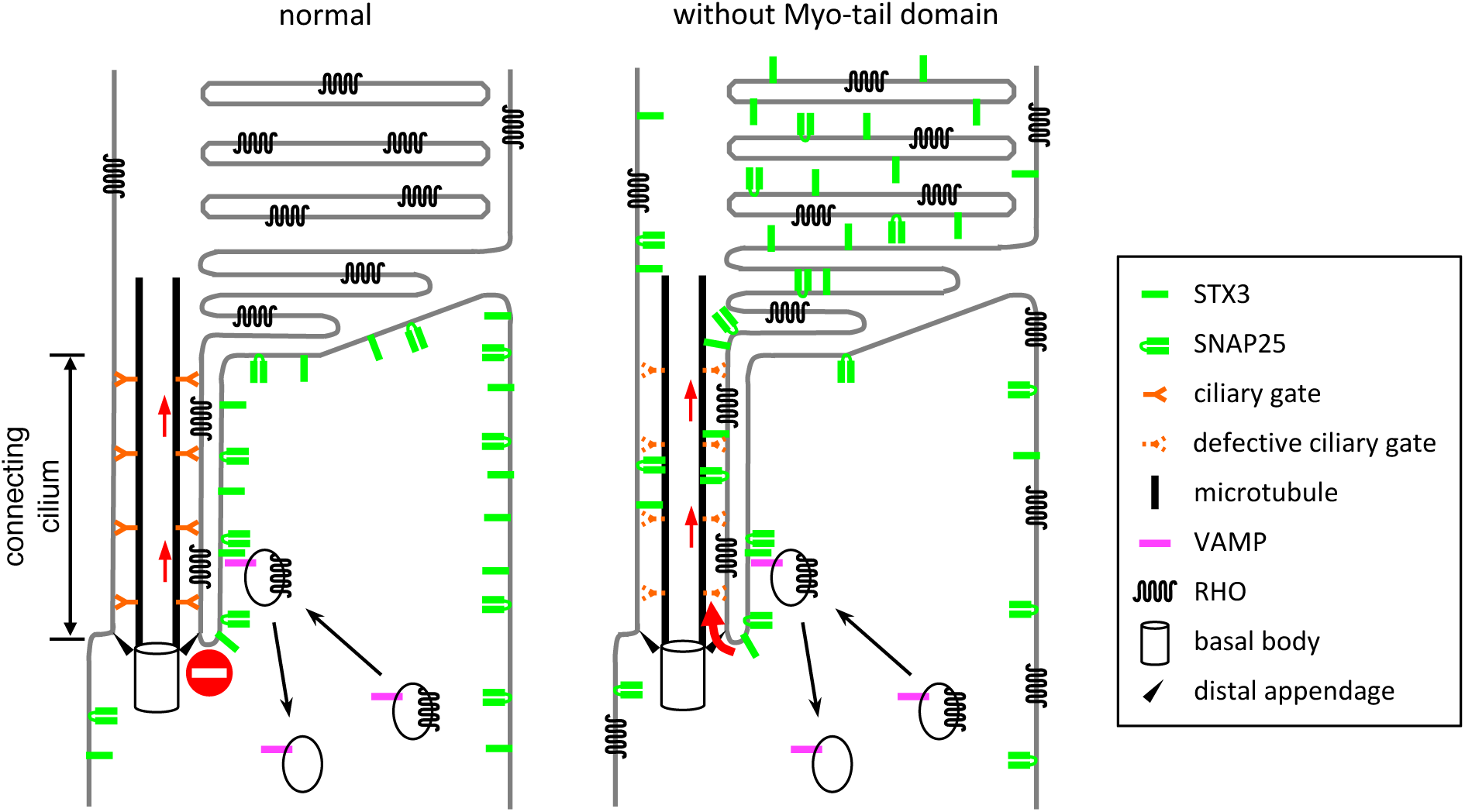
A model for pathomechanisms of *CEP290*-associated retinopathies. CEP290 is a component of the Y-link that functions as a ciliary gatekeeper at the connecting cilium. In normal photoreceptors (*left*), CEP290 allows entry of outer segment-bound proteins while blocking that of inner segment proteins such as STX3, STXBP1, SNAP25, and IMPG2. In photoreceptors with compromised CEP290 function (*right*), defective ciliary gates allow entry of not only outer segment-bound proteins but also certain inner segment membrane proteins that do not have specific targeting signals. Although precise roles of CEP290 in RHO trafficking are currently uncertain, a subset of RHO mislocalizes to the inner segment and RHO mislocalization is likely to contribute to retinal degeneration in *CEP290*-associated ciliopathies.

## MATERIALS AND METHODS

### Animal study approval

All animal procedures were approved by the Institutional Animal Care and Use Committee (IACUC) of the University of Iowa and conducted in accordance with the recommendations in the Guide for the Care and Use of Laboratory Animals of the National Institutes of Health.

### Mouse

*Cep290*^*fl*^ (Cep290^tm1Jgg^/J; stock number 013701) [26] and *Cep290*^*rd16*^ (BXD24/TyJ-Cep290^rd16^/J; stock number 000031) mice were obtained from the Jackson laboratory. *iCre75* transgenic mice [27] were obtained from Dr. Ching-Kang Chen (Baylor College of Medicine). *Ift88*^*fl*^ and *Lztfl1* gene-trap mice were from our colonies and previously described [45, 70]. The original *Cep290*^*fl*^ mice that we obtained had both *rd1* (in *Pde6b*) and *rd8* (in *Crb1*) mutations. These mutations were eliminated by breeding with wild-type 129S6/SvEvTac mice (Taconic Biosciences), and all animals used in this study were *Pde6b*^*+/+*^*;Crb1*^*+/+*^ in B6;129S6 mixed backgrounds. *Cep290*^*rd16*^ mice were maintained in a BXD;B6 mixed background to obtain heterozygous littermates (i.e. *Cep290*^*+/rd16*^) as a control. All primers for genotyping were from Integrated DNA Technologies and their sequences are described in **Table 1**. All animals were maintained in 12-hour light/dark cycles and fed *ad libitum* standard mouse chow. PCR protocols are available upon request.

**Table 1.**
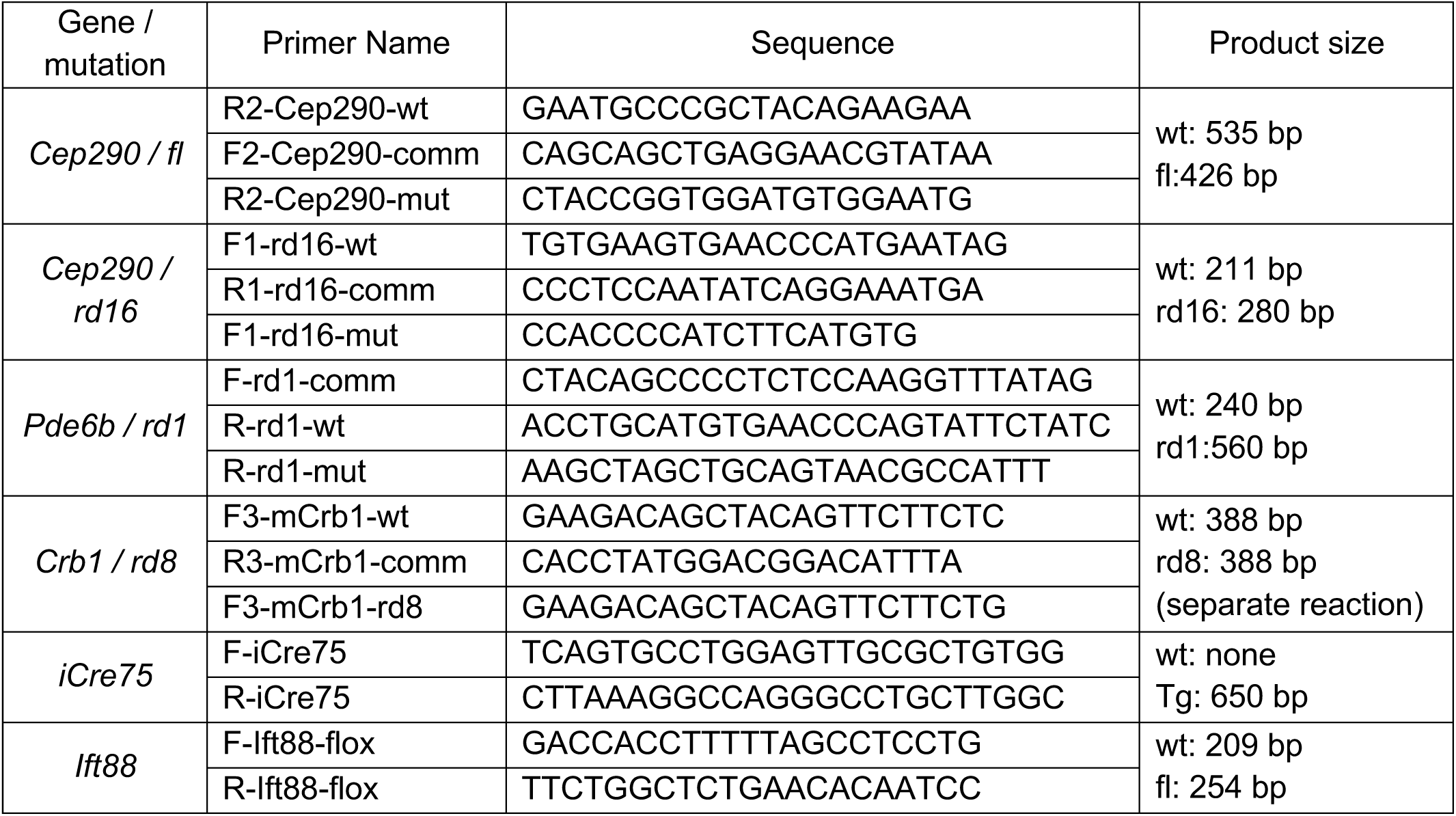
PCR primers used for genotyping.

### RNA extraction and Reverse Transcription (RT)-PCR

Mice were euthanized by CO_2_ asphyxiation followed by cervical dislocation. Eyes were enucleated and the anterior segment was removed using micro-dissecting scissors. The neural retina was separated from the rest of the ocular tissues with forceps and snap-frozen in liquid nitrogen. Upon completion of retina collection, frozen retinas were placed on ice, immersed in 1 ml of TRI Reagent (Sigma), and homogenized with PT1200E Polytron homogenizer (Kinematica). Total RNAs were extracted following the manufacturer’s instruction. One μg of total RNA was used for cDNA synthesis using SuperScript IV Reverse Transcriptase (Thermo Fisher Scientific) and random hexamers (Thermo Fisher Scientific) following the manufacturer’s instruction. *Cep290* cDNA fragments between exons 34 and 41 were PCR amplified with Universe High-Fidelity Hot Start DNA polymerase (Bimake) and two primers (F-Cep290-e34: 5’-GCCGAAATCTCATCACACAATG-3’ and R-Cep290-e41: 5’-GCTTCTCCTTCCTTCTCCTTTAG-3’). PCR products were separated in a 1.2% agarose gel and purified with a Gel/PCR DNA Fragment Extraction kit (IBI Scientific). Purified DNAs were sequenced by Sanger sequencing using the aforementioned two primers (F-Cep290-e34 and R-Cep290-e41).

### Immunofluorescence microscopy

After euthanasia by CO_2_ asphyxiation followed by cervical dislocation, mouse eyes were enucleated and immersed in 4% (wt/vol) paraformaldehyde (PFA)/PBS fixative. A small puncture was created between the lens and the sclera using a 26-G needle. After 5 minutes of fixation, the lens and the anterior chamber were removed with micro-dissecting scissors. The eye-cups were further fixed in 4% PFA/PBS for 3 hours at 4 °C. After washing with PBS (3 times, 10-min each), eye-cups were infiltrated and embedded in acrylamide as previously described [71]. After solidification, extra acrylamide was carefully removed and eye-cups were placed in Neg-50 Frozen Section Medium (Thermo Fisher Scientific), frozen in a dry-ice/ethanol bath, and stored at −80 °C. Five to eight μm cryosections were collected on Superfrost Plus Microscope Slides (Fisher Scientific) using a CryoStar NX70 Cryostat (Thermo Fisher Scientific). Retinal sections were permeabilized with PBS-T (PBS with 0.1% Triton X-100), blocked with 5% BSA/5% normal goat serum (Vector Laboratories) in PBS-T, and incubated with indicated primary antibodies for 2-3 hours at room temperature. After washing, sections were incubated with secondary antibodies conjugated to Alexa fluor 488 or 568 (Thermo Fisher Scientific) for 2 hours at room temperature in the dark. After rinsing, Vectashield Mounting Medium with 4′,6-Diamidine-2′-phenylindole dihydrochloride (DAPI) (Vector Laboratories) was added and images were taken using Olympus IX71 microscope or Zeiss LSM 710 confocal microscope. Intensity range of images was adjusted by linear level adjustments using Adobe Photoshop CC 2018 (Adobe Systems). For fluorescence intensity quantification, 20-25 μm width areas were randomly selected and integrated fluorescence intensities were measured within the outer segment and the entire photoreceptor cells using ImageJ to calculate the ratio of fluorescence intensities in the outer segment. Two-tailed, two-sample *t*-tests assuming unequal variances were used for statistical analyses. *P* values smaller than 0.01 were regarded as statistically significant.

### Outer segment isolation and immunoblotting

Photoreceptor outer segments were isolated as previously described [45]. Briefly, eyes were enucleated from *Cep290*^*+/fl*^*;Cre*^*?*^ and *Cep290*^*fl/fl*^*;Cre*^*+*^ mice at P25-28, and the anterior segment and the lens were removed using micro-dissecting scissors. The neural retina was separated from the pigmented retinal epithelium with forceps, collected in 63% sucrose/PBS, snap-frozen in liquid nitrogen, and stored in −80 °C until needed. Fifteen to twenty animals were used per genotype per preparation. Retinas with same genotypes were pooled in 1.5-ml tubes on ice and 63% sucrose/PBS was added to 1 ml. Retinal suspensions were gently pipetted up and down 10 times using a P1000 tip with a 1.5-2 mm orifice, and further vortexed for 30 seconds. Homogenates were centrifuged at 100 × g for 3 minutes, and supernatants were transferred to fresh 1.5-ml tubes and spun at 2,350 × g for 10 minutes at 4 °C. Supernatants were transferred to ultracentrifuge tubes, and layers of 42%, 37%, and 32% sucrose/PBS solutions (1.1 ml each) were overlaid. Homogenates were centrifuged at 116,000 × g_ave_ for 1 hour at 4°C using a Sorvall TH-660 rotor. Outer segments were collected at the 32%-37% sucrose interface, and an equal volume of ice-cold PBS was added to the collected outer segment fractions. Outer segments were pelleted by spinning at 10,000 × g for 8 minutes at 4 °C and resuspended in a lysis buffer (PBS with 1% Triton X-100). Protein concentrations were measured with a DC Protein Assay kit (Bio-Rad) following the manufacturer’s instruction. Five μg of proteins were loaded onto a 4-12% (wt/vol) NuPAGE Bis-Tris gel (Thermo Fisher Scientific), and SDS-PAGE and immunoblotting were conducted following standard protocols. Proteins were detected with SuperSignal West Dura Extended Duration Substrate (Thermo Fisher Sciencetific). Images were taken with a ChemiDoc system (Bio-Rad) and quantified with the Image Lab software (Bio-Rad).

### Immunoprecipitation

Neural retinas from *Cep290*^*+/fl*^*;Cre*^*?*^ and *Cep290*^*fl/fl*^*;Cre*^*+*^ mice were isolated as described above and lysed in 400 μl of a lysis buffer (50 mM HEPES pH 7.0, 150 mM NaCl, 0.5% NP-40, 2 mM EGTA, 2 mM MgCl_2_, 10% glycerol, protease inhibitor cocktail (Bimake)) by pipetting and vortexing. After removing insoluble materials by centrifugation at 15,000 x g for 10 minutes, 1.5-2 μg of BBS7 (Proteintech Group; 18961-1-AP), STX3 (Proteintech Group; 15556-1-AP), or rabbit polyclonal control IgG (Proteintech Group; 30000-0-AP) antibodies were added to supernatants and incubated at 4 °C overnight with rotation. To precipitate antibodies and associated proteins, 10 μl of Dynabeads Protein G magnetic beads (Thermo Fisher Scientific) were added and further incubated at 4 °C for 2 hours with rotation. Beads were washed 4 times with 1 ml of a wash buffer (50 mM HEPES pH 7.0, 150 mM NaCl, 0.1% NP-40, 2 mM EGTA, 2 mM MgCl_2_, 10% glycerol). Proteins were eluted by adding 25 μl of 1x NuPAGE LDS sample buffer (Thermo Fisher Scientific) and 20 μl was loaded onto a 4-12% (wt/vol) NuPAGE Bis-Tris gel (Thermo Fisher Scientific). SDS-PAGE, immunoblotting, and imaging were conducted as described above except that rabbit primary antibodies were detected by HRP-conjugated Protein A (Proteintech Group; SA00001-18).

### Antibodies

The following primary antibodies were used for immunofluorescence microscopy and immunoblotting: ABCA4 (EMD Millipore; MABN2439), β-actin (Sigma; A1978), ATP1A3 (abcam; ab2826), BBS2 (Proteintech Group; 66246-1-AP), BBS5 (Santa Cruz Biotechnology; sc-515331), BBS7 (Proteintech Group; 18961-1-AP), BBS8 (Sigma; HPA003310), CEP290 (EMD Millipore; ABN1710), CETN (EMD Millipore; 04-1624), CNGA1 (EMD Millipore; MABN468), FLAG (Sigma; F1804), GRK1 (abcam; ab2776), GUCY2D (Proteintech Group; 55127-1-AP), HCN1 (NeuroMab; 75-110), IMPG2 (Thermo Fisher Scientific; PA5-64926), LAMP1 (Sigma; L1418), LZTFL1 (custom-made; [66]), PDE6A (Proteintech Group; 21200-1-AP), PDE6B (Proteintech Group; 22063-1-AP), PRPH2 (Proteintech Group; 18109-1-AP), RHO (EMD Millipore; MAB5356), ROM1 (Proteintech Group; 21984-1-AP), SNAP25 (abcam; ab24737), STX3 (EMD Millipore; MAB2258), STXBP1 (Proteintech Group; 11459-1-AP), SYP (Cell Signaling; 5461) and VAMP2 (Cell Signaling; 13508). Rabbit polyclonal ATP8A2 antibody was custom-made using GST-tagged mouse ATP8A2 (amino acids 369-644) as an immunogen, followed by affinity-purification with MBP-tagged mouse ATP8A2 (amino acids 369-644) (Proteintech Group).

## ACKNOWLEDGEMENTS

We thank Dr. Ching-Kang Chen (Baylor College of Medicine) for generously providing *iCre75* transgenic mice. This work was supported by National Institutes of Health grants R01-EY022616 and R21-EY027431 (to S.S.) and National Institutes of Health Center Support grant P30-EY025580 to the University of Iowa.

