## Supplemental information text for "CEP290 myosin-tail homology domain is essential for protein confinement between inner and outer segments in photoreceptors"

**Supplementary Figure Legends**

**Supplementary Figure S1**. **Connecting cilium assembly and the initial development of outer segments in *Cep290$^{fl/fl}$;Cre$^{+}$* retinas is not disrupted until PN11.**

Retinal sections from 11-days old *Cep290$^{+/fl}$;Cre$^{+}$* and *Cep290$^{fl/fl}$;Cre$^{+}$* mice were immunostained with indicated antibodies. (A) Connecting cilia and basal bodies are labeled with anti-CETN antibody (red). CEP290 (green) is detected at the connecting cilium in both *Cep290$^{+/fl}$;Cre$^{+}$* and *Cep290$^{fl/fl}$;Cre$^{+}$* mice until PN11. (B-D) Initial development of the outer segment appears to be unaltered in *Cep290$^{fl/fl}$;Cre$^{+}$* retinas until PN11. Note that mild mislocalization of RHO is observed in both *Cep290$^{+/fl}$;Cre$^{+}$* and *Cep290$^{fl/fl}$;Cre$^{+}$* retinas until this age and that localization of PRPH2 and ROM1 is also comparable. Nuclei were counterstained with DAPI (blue). Scale bars represent 10 $\mu$m.

**Supplementary Figure S2. Cleavage and localization of IMPG2 in HEK293T cells.**

(A) Cleavage of IMPG2. HEK293T cells were transfected with a construct to express mouse *Impg2* (NM_174876) with a 2xS and 3xFLAG (SF) tag at the C-terminus. Mock-transfected cells were used as a negative control. Protein extracts were subjected to SDS-PAGE and immunoblotting with anti-IMPG2 and anti-FLAG antibodies. Although calculated molecular weight of full-length mouse IMPG2 (without glycosylation and modification) is 138 kDa and that of the SF tag is 13 kDa, observed molecular weights of the two major bands (black arrowheads) detected by anti-IMPG2 and anti-FLAG antibodies are 40-45 kDa (with the SF tag), indicating that full-length IMPG2 is rapidly cleaved in HEK293T cells. Red arrowhead indicates presumptive full-length IMPG2 detected by anti-FLAG antibody after over-exposure. Asterisk denotes a cross-reacting protein detected by anti-IMPG2 antibody. Anti-IMPG2 antibody was raised to the C-terminal intracellular domain of human IMPG2 (aa 1127-1234).

(B) Plasma membrane localization of the IMPG2 C-terminal cleavage products. HEK293T cells were transfected with the IMPG2-SF expression cassette described in (A), and localization of IMPG2-SF was probed with anti-IMPG2 (green) and anti-FLAG (red) antibodies. Note that the vast majority of IMPG2-

SF detected by these antibodies are C-terminal cleavage products (shown in A; black arrowheads), not full-length proteins or N-terminal fragments. White arrowheads indicate plasma membrane localization and open arrowheads denote nuclei of untransfected cells. DAPI was used to counterstain nuclei (blue).

**Supplementary Figure S3. Expression and localization of STX3 in cone synapses.**

Retinal sections from 40-days old *Cep290$^{fl/fl}$;Cre$^{+}$* mice were stained with STX3 antibodies (green) and a cone cell marker, peanut agglutinin (PNA; red). Note that STX3 staining in the synaptic terminal layer overlaps with PNA (red arrowheads), indicating that STX3 in the synaptic terminal layer is mostly from cones, in which *Cre* is not expressed. White dashed lines mark an area where cones are absent. Note the depletion of STX3 in the synaptic terminal layer. Open arrowhead denotes a blood vessel.

**Supplementary Figure S4. Co-immunoprecipitation of STX3 and STXBP1.**

Retinal protein extracts from wild-type mice were subjected to immunoprecipitation (IP) with anti-STX3 antibody. Normal rabbit polyclonal IgG was used as a negative control. Precipitated proteins were separated by SDS-PAGE and visualized by silver staining. While STX3 interactions with SNAP25 and VAMP2 are detectable by immunoblotting, they are not visible in a silver stained gel. In contrast, STX3-STXBP1 interaction is readily detectable by silver staining with a near-stoichiometric proportion.
