## Supplemental information figures for "CEP290 myosin-tail homology domain is essential for protein confinement between inner and outer segments in photoreceptors"

### Supplementary Figure S1

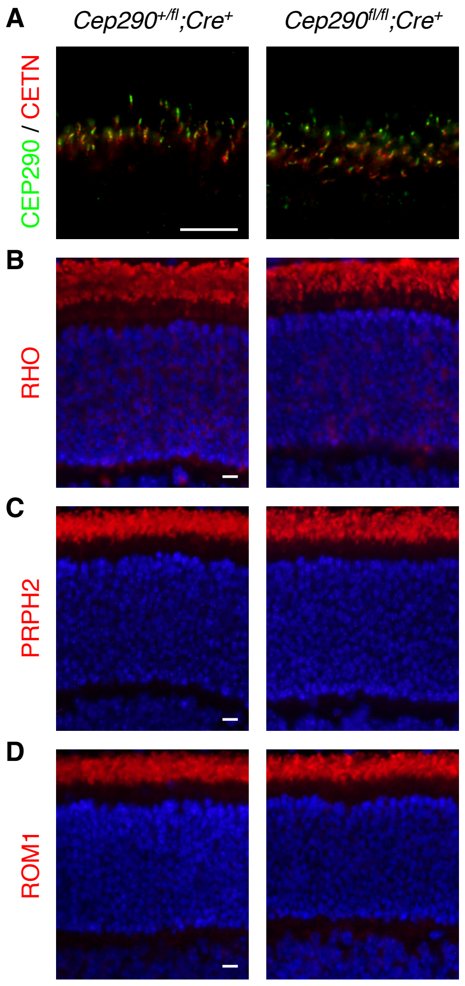

### Supplementary Figure S2

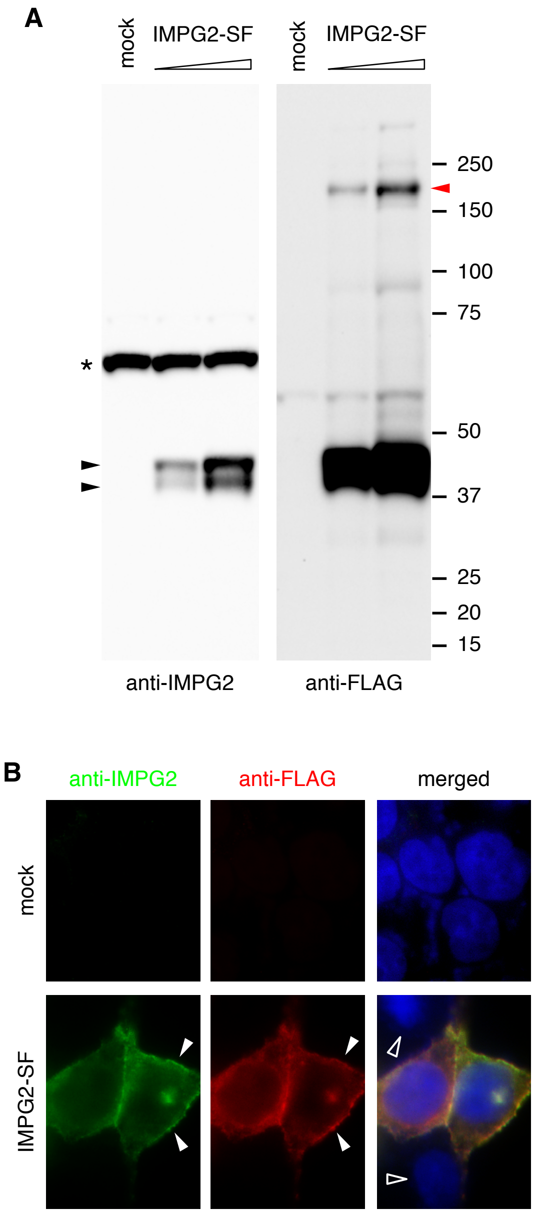

**Supplementary Figure S3**

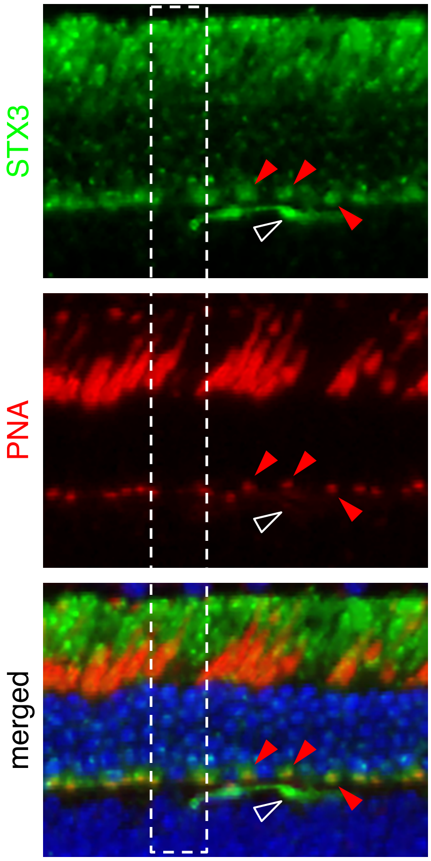

#### Supplementary Figure S4

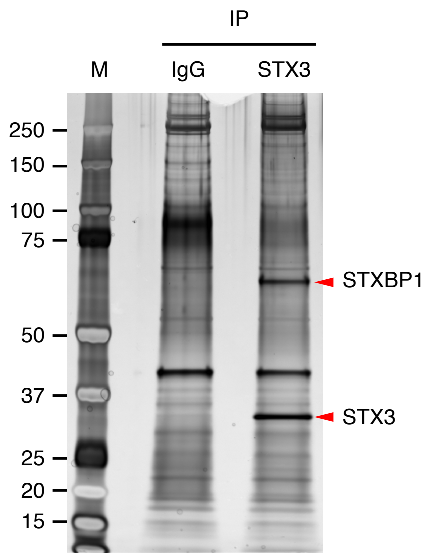
